# Actomyosin and the Arp2/3 Complex Are Involved in the Internalization of Cellulose Synthase Complexes

**DOI:** 10.1101/2025.01.19.633783

**Authors:** Liyuan Xu, Weiwei Zhang, Lei Huang, Chunhua Zhang, Christopher J. Staiger

**Affiliations:** Department of Biological Sciences, Purdue University, West Lafayette, IN 47907, USA; Department of Botany and Plant Pathology, Purdue University, West Lafayette, IN 47907, USA; EMBRIO Institute, Purdue University, West Lafayette, IN 47907, USA

**Keywords:** actin cytoskeleton, Arp2/3 complex, myosin, cellulose synthase complex, exocytosis, endocytosis, internalization, photoconversion, microscopy

## Abstract

The coupling of exo- and endocytic trafficking of Cellulose Synthase Complexes (CSCs) has been proposed to be important for maintaining the population of active CSCs at the plasma membrane (PM) and thus appropriate levels of cell wall assembly. Although actin and myosin are known to participate in the late stages of exocytosis of CSCs, their exact role during CSC internalization events remains controversial. We constructed a functional, photoconvertible fluorescent mEOS2-CESA6 reporter and developed single-particle live-cell imaging approaches to visualize and quantify the dynamic behavior of CSCs at the PM during internalization. Using the small molecule inhibitor of clathrin, Endosidin 9-17 or ES9-17, we confirmed that clathrin-mediated endocytosis is a major pathway for CSC internalization. We also found that the actin cytoskeleton is involved in CSC internalization. Genetic or chemical inhibition of actin, myosin, or the Arp2/3 complex significantly reduced the frequency of CSC internalization events and prolonged the CSC pause time prior to internalization. Additionally, we found that the Arp2/3 complex contributes to the late stage of exocytosis of CSCs into the PM. These results reveal a role for actomyosin and the Arp2/3 complex in both CSC secretion as well as internalization that was previously undescribed in plant cells.

**One sentence summary:** Direct visualization of individual CSC internalization events reveals that actomyosin participates in CSC internalization and the Arp2/3 complex contributes to both exocytosis and internalization of CSCs through regulating the dynamic homeostasis of the cortical actin cytoskeleton.

## Introduction

Exocytosis and endocytosis are two fundamental cellular processes. The coupling of exocytic and endocytic trafficking plays an essential role in regulating multiple cell functions, including signal transduction, cell polarity establishment, material recycling, as well as in modulating the homeostatic level of plasma membrane (PM)-proteins (Wu et al., 2014; Fan et al., 2015; Kaksonen and Roux, 2018; Zhang et al., 2019; Zhu and McFarlane, 2022). In yeast and animal cells, the actin cytoskeleton is a highly dynamic system that undergoes rapid rearrangement, and it cooperates with myosin for both exo- and endocytic trafficking in response to biotic and abiotic stimuli (Porat-Shliom et al., 2013; Miklavc and Frick, 2020; Wu and Chan, 2022). In plant cells, actin and myosin are responsible for long distance intracellular trafficking and cytoplasmic streaming (Reddy and Day, 2001; Beck et al., 2012; Wang and Hussey, 2015; Breuer et al., 2017). Actomyosin also participates in the late stages of exocytosis (Sampathkumar et al., 2013; Zhang et al., 2019; Zhang et al., 2021; Zhang and Staiger, 2021; Liu et al., 2023), but its contribution to endocytosis remains controversial.

The Cellulose Synthase (CESA) Complex (CSC) is an excellent cargo marker for studying plant intracellular trafficking as well as material secretion and/or internalization. Expressing a functional fluorescence-tagged CESA6 in the *cesa6^prc1^* mutant background allows us to visualize the trafficking of individual CSC particles to and from the PM (Paredez et al., 2006; Gutierrez et al., 2009; Watanabe et al., 2015). CSCs are first assembled in the Golgi (Giddings et al., 1980; Haigler and Brown, 1986; Paredez et al., 2006), and they are packaged into a subpopulation of small cytoplasmic CESA compartments (SmaCCs) (Gutierrez et al., 2009) or microtubule-associated transport vesicles (MASCs) (Crowell et al., 2009). Then, they are delivered to the cortical cytoplasmic region through post-Golgi CESA trafficking (Zhu and McFarlane, 2022). After successful insertion into the PM, CSCs get activated and start producing cellulose microfibrils on the cell surface (Mueller and Brown, 1980; Arioli et al., 1998; Somerville, 2006; Sampathkumar et al., 2013; Zhu and McFarlane, 2022). The actin cytoskeleton and myosins XI play a key role during the late stages of CSC exocytosis, specifically the tethering, docking, and fusion of CSC exocytic vesicles to the PM (Sampathkumar et al., 2013; Zhang et al., 2019; Zhang et al., 2021; Zhang and Staiger, 2021). In addition, myosin XIK, the primary isoform of plant myosin XI, interacts directly with the exocyst complex subunits, Sec5B and Sec15, to facilitate the tethering of CSC exocytic vesicles to the PM (Zhang et al., 2019; Zhang et al., 2021). A recent study also shows that actin and myosin XI are involved in a non-canonical pathway, in which actomyosin participates in the tail-stretching and breaking of Golgi membranes so that CSCs are delivered directly from Golgi to the PM (Liu et al., 2023). Surprisingly, although the delivery of new CSCs to the PM is reduced when actin is perturbed genetically or chemically, the abundance of functional CSCs at the PM is unaltered or only modestly reduced (Sampathkumar et al., 2013; Zhang et al., 2019). This implies that the actin cytoskeleton plays an equal and opposite role during CSC secretion and internalization.

In contrast to exocytosis, the endocytic process of CSCs has not been dissected with near the same detail (Zhu and McFarlane, 2022). Disrupting the activity of several key players of plant clathrin-mediated endocytosis (CME), including adaptor proteins, dynamin-related proteins, and clathrin proteins, all lead to increased density of PM-localized CSCs but with significantly reduced motility, suggesting that CME may participate in internalization of dysfunctional or inactive CSCs (Bashline et al., 2013; Bashline et al., 2015; Sánchez-Rodríguez et al., 2018). After internalization, CSC endocytic vesicles are transferred into the cytoplasm through early endosome/trans-Golgi network (EE/TGN), which bring CSCs back to the PM through recycling or to the vacuole for degradation (Endler et al., 2015; Lei et al., 2015; Sánchez-Rodríguez et al., 2018; McFarlane et al., 2021; Zhu and McFarlane, 2022). Several studies show that recycling plays an important role in maintaining the activity of CSCs, because recycling has been proposed as a much faster way to recover large functional protein complexes compared to new synthesis (Ivanov and Vert, 2021; McFarlane et al., 2021; Zhu and McFarlane, 2022). However, it remains to be determined whether CSC recycling occurs through CME or through clathrin-independent pathways. The coordination between secretion, recycling, and internalization of CSCs likely determines the abundance of active CSCs at the PM and consequently affects cellulose synthesis and cell wall assembly (Bashline et al., 2014; Wallace and Somerville, 2014). However, no study directly visualizes and quantifies the stages of CSC internalization; thus, it is unclear whether actin or myosin are involved in this process. Previous studies show conflicting results regarding the involvement of actin cytoskeleton in plant endocytosis, either the constitutive endocytosis of FM4-64 lipid membrane dye (Moscatelli et al., 2012; Sampathkumar et al., 2013; Kang et al., 2014; Mao et al., 2016; Zhang et al., 2019; Narasimhan et al., 2020) or the ligand-triggered endocytosis of FLS2 (Robatzek et al., 2006; Beck et al., 2012; Zhang et al., 2019). A recent study uses scanning electron microscopy to directly visualize the clathrin coat/basket *in situ*, observing only thick actin bundles but not the expected dense actin meshwork surrounding endocytic foci near the PM (Narasimhan et al., 2020).

The Actin-related Protein 2/3 complex (Arp2/3), a major nucleator of side-branched actin filaments (Machesky et al., 1994; Mullins et al., 1998; Winter et al., 1999; Amann and Pollard, 2001; Robinson et al., 2001), is critical for plant cells to maintain the dynamic homeostasis of the cortical actin network through cooperating with formins to regulate the actin assembly (Xu et al., 2024). The Arp2/3 complex is also important for cell wall construction and epidermal cell morphology development in Arabidopsis (Le et al., 2003; Li et al., 2003; Mathur et al., 2003; Sahi et al., 2018; Liu et al., 2023; Xu et al., 2024). In addition, *arp2/3* mutants show vacuole morphology defects, suggesting that Arp2/3 is essential for vacuolar fusion or even general membrane fusion (Eitzen, 2003; García-González et al., 2020). Furthermore, several recent studies found that AtEH/Pan1, a subunit of TPLATE COMPLEX (TPC, an plant-specific adaptor complex)(Gadeyne et al., 2014), binds to the actin cytoskeleton and recruits the Arp2/3 complex and two other TPC subunits, TPLATE and TML, during the formation of autophagosomes, indicating the direct interaction between the Arp2/3 complex and plant endocytic machinery (Wang et al., 2016; Wang et al., 2019; Wang et al., 2020). Collectively, these data suggest that the Arp2/3 complex and Arp2/3-mediated actin networks may be essential important for cellulose synthesis, which could be accomplished through participating in both exo- and endocytosis of CSCs. Direct evidence for this hypothesis is currently lacking.

In this study, we developed a quantitative imaging approach to directly visualize and assess CSC internalization by tracking individual CSC particles during their disappearance from the PM, using a new photoconvertible mEOS2-CESA6 marker. We found that actin, the Arp2/3 complex, and myosin all participate in CSC internalization, possibly through both clathrin-mediated and clathrin-independent endocytosis. Actomyosin and the Arp2/3 complex contribute to the coupling of exo- and endocytic trafficking of CSCs and maintain the population of active CSCs on the PM, thereby playing a role in cellulose production and cell wall assembly.

## Results

### Cellulose levels are reduced in *arp2/3* mutants of Arabidopsis

Arabidopsis *arp2/3* mutants show pronounced growth reduction in both dark-grown hypocotyls and light-grown roots as well as exhibit severe defects in epidermal cell morphology (Le et al., 2003; Li et al., 2003; Mathur et al., 2003; El-Assal et al., 2004; Sahi et al., 2018; Xu et al., 2024). To test whether the Arp2/3 complex is involved in cellulose production, we measured total and crystalline cellulose content in alcohol-insoluble cell wall fractions prepared from 5-day-old etiolated hypocotyls of homozygous *arp2/3* mutants (*arp2-1*, *arp2-2*, and *arpc2*), the *cesa6/prc1-1* mutant, as well as wild-type sibling lines. Both total and crystalline cellulose content in *arp2/3* mutant hypocotyls were significantly lower than in the wild type, however, were not as severe as the *prc1-1* mutant, suggesting that the Arp2/3 complex participates in cellulose biosynthesis (Supplemental Fig. S1).

### The Arp2/3 complex is involved in the late stages of exocytosis of Cellulose Synthase Complexes (CSCs) to the plasma membrane (PM)

Previous studies demonstrate that both actin and myosins XI are involved in the late stages of CSC exocytosis; however, inhibition of actin genetically or chemically does not alter the abundance of CSCs at the PM, suggesting that the actin cytoskeleton might participate in both secretion and internalization of CSCs (Sampathkumar et al., 2013; Zhang et al., 2019; Zhang et al., 2021). We confirmed these findings using treatment with the actin polymerization inhibitor Latrunculin B (LatB) as well as the myosin inhibitor pentabromopseudilin (PBP; Fig. 1). To test whether the Arp2/3 complex plays a similar role during the intracellular trafficking of CSCs, we employed genetic and chemical inhibitor approaches. We crossed the *arp2-2* and *arpc2* mutants with a line expressing YFP-CESA6 in the *cesa6*/*prc1-1* mutant background (Paredez et al., 2006), and recovered both *arp2-2* and *arpc2* homozygous mutants as well as wild-type sibling lines expressing YFP-CESA6 in the *cesa6*/*prc1-1* background. In addition, we also used a small-molecule inhibitor of the Arp2/3 complex, CK-666 (Hetrick et al., 2013), for acute inhibition of actin nucleation activity of the Arp2/3 complex (Xu et al., 2024).

**Figure 1.**
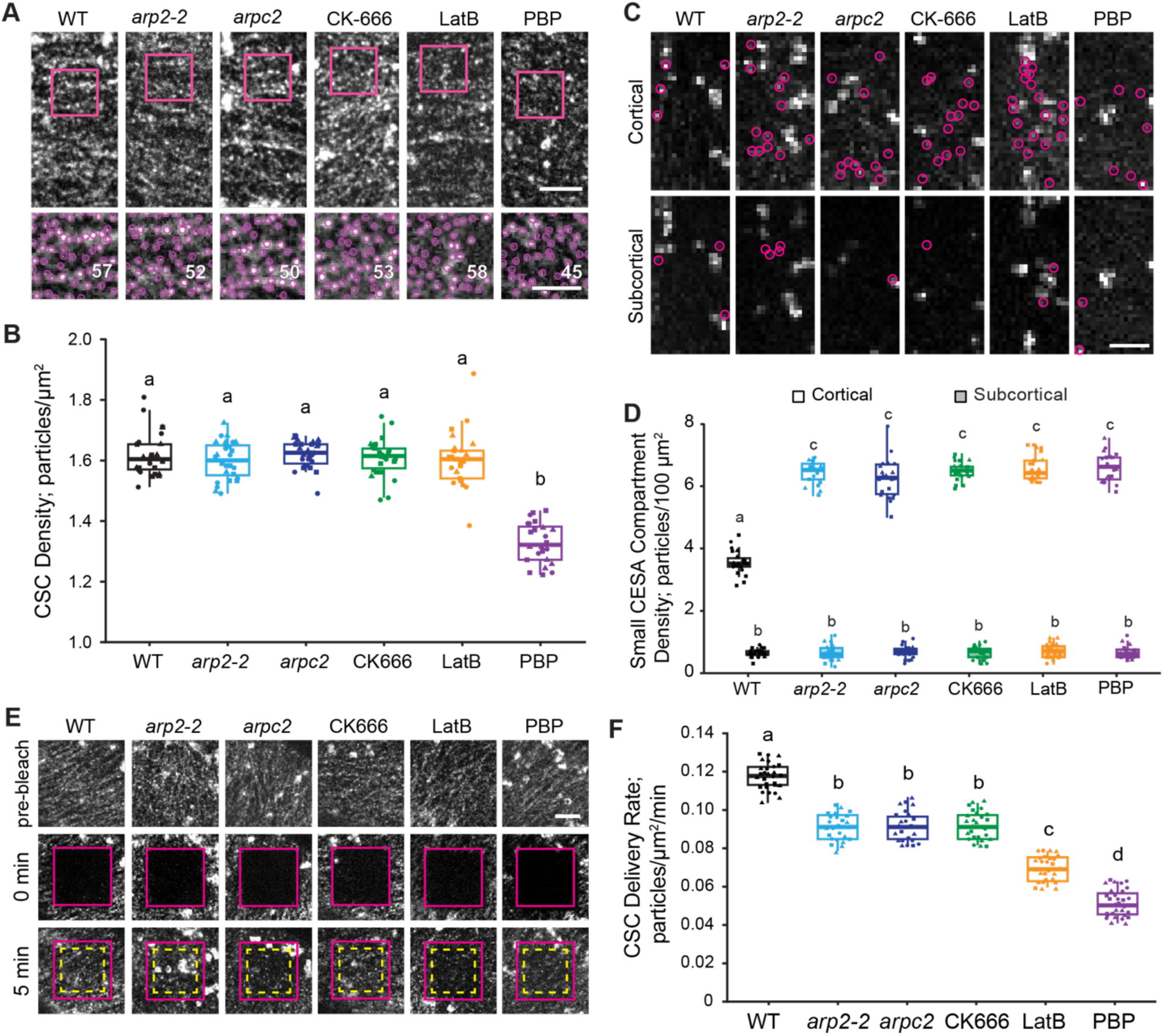
Inhibition of the Arp2/3 complex reduces the delivery of Cellulose Synthase Complexes (CSCs) into the plasma membrane (PM). A, Representative single-frame images of epidermal cells from the apical region of 3-d-old etiolated hypocotyls expressing YFP-CESA6 collected by spinning disk confocal microscopy (SDCM) show the distribution of CSCs in the PM (top row). Bar = 5 μm. The number of CSC particles detected with TrackMate in ImageJ (circles) in a region of interest or ROI (box) is shown in the bottom row. Bar = 3 μm. Seedlings from wild-type Col-0 lines (WT) were treated with mock solution (0.05% DMSO), 10 µM CK-666, 10 µM LatB, or 10 µM PBP for 15 min prior to imaging with SDCM. Seedlings of *arp2-2* and *arpc2* were mock-treated with 0.05% DMSO for 15 min. B, Quantitative analysis of CSC abundance at the PM. The density of CSCs was not significantly different in *arp2-2*, *arpc2*, CK-666-treated, or LatB-treated cells when compared to mock-treated wild-type cells. However, PBP treatment significantly reduced the density of CSCs in the PM compared to mock treatment. C, Representative single-frame images of hypocotyl epidermal cells taken at the cortical (top) or subcortical (bottom) focal plane show the distribution of cytoplasmic small CESA compartments (SmaCCs; circles). Bar = 5 μm. D, Quantitative analysis of SmaCC abundance in the cortical or subcortical focal plane. The number of SmaCCs in the cortical cytoplasm was significantly increased in *arp2-2*, *arpc2*, CK- 666-, LatB-, and PBP-treated cells when compared to that in mock-treated wild-type cells. In contrast, SmaCC density in the subcortical cytoplasmic region did not show a significant difference between mock-treated wild type and any mutant or chemical inhibitor treatment. E, Representative single-frame images illustrate the fluorescence recovery of PM-localized CSC particles following photobleaching. Genotypes and treatments were as described above. An ROI at the PM was photobleached (magenta boxes; middle row) and the number of newly-delivered CSCs was counted in a subarea within the photobleached region (yellow dashed boxes; bottom row). Bar = 5 μm. F, Quantitative analysis of the rate of delivery of functional CSCs to the PM. The delivery rate was calculated from the total number of newly-delivered CSCs identified during the first 5 min of recovery after photobleaching divided by the measured area and time. The CSC delivery rate was significantly reduced in *arp2-2*, *arpc2*, or CK-666-treated cells when compared to mock-treated wild-type cells; however, these reductions were not as severe when compared to the reduction observed for PBP- or LatB-treated cells. In box-and-whisker plots, boxes show the interquartile range (IQR) and the median, and whiskers show the maximum-minimum interval of three biological repeats with independent populations of plants. The three independent experiments are represented with different symbols. n ≥ 21 seedlings (at least 7 seedlings were measured per independent experiment). Letters [a–d] denote genotypes or treatments that show statistically significant differences with other groups by one-way ANOVA with Tukey’s post-hoc test (P < 0.05).

To test whether the reduced cellulose content in *arp2/3* mutants correlates with reduced abundance of CSCs in the PM, we quantified the density of CSCs at the PM by comparing images of epidermal cells from the apical region of 3-day-old wild-type and Arp2/3-inhibited etiolated hypocotyls collected with high spatiotemporal resolution spinning disk confocal microscopy (SDCM; Fig. 1A; Gutierrez et al., 2009; Sampathkumar et al., 2013; Zhang et al., 2019; Zhang et al., 2021). We found that the density of PM-localized CSCs in *arp2-2*, *arpc2*, or CK-666-treated cells was not significantly different compared to mock-treated wild-type cells (Fig. 1B). The catalytic activity of CESA correlates with the speed of linear motility of CSCs in the PM (Paredez et al., 2006; Huang et al., 2023). To test whether the cellulose synthesis activity of CSCs was altered in *arp2/3* mutants, we measured the motility of CSCs in the plane of PM, as described previously (Supplemental Fig. S2, A and B; Paredez et al., 2006; Zhang et al., 2019; Huang et al., 2023). The inhibition of the Arp2/3 complex caused a minor reduction in the average CSC speed of CSC motility in the plane of the PM (Supplemental Fig. S2C).

To investigate whether the Arp2/3 complex is required for intracellular trafficking of CSCs, we quantified the abundance of intracellular small CESA compartments (SmaCCs) in the cortical (0–0.4 μm below the PM) and subcortical cytoplasm (0.6–1.0 μm below the PM; Fig. 1C; Gutierrez et al., 2009; Sampathkumar et al., 2013; Zhang et al., 2019; Zhang et al., 2021).

Defects in the late stages of exocytosis lead to an accumulation of cortical SmaCCs, a subpopulation of which are thought to represent secretory vesicles (Sampathkumar et al., 2013; Zhang et al., 2019). Compared to mock-treated wild-type cells, both genetic and chemical inhibition of the Arp2/3 complex caused a significant accumulation of SmaCCs in the cortical cytoplasmic region (Fig. 1D). However, the amount of SmaCCs in the subcortical cytoplasm did not show any significant differences between genotypes or treatments (Fig. 1D). To determine whether the Arp2/3 complex contributes to the delivery of functional CSCs into the PM, we performed fluorescence recovery after photobleaching (FRAP) experiments, as described previously (Fig. 1E; Gutierrez et al., 2009; Bashline et al., 2013; Sampathkumar et al., 2013; Luo et al., 2015; Zhang et al., 2019; Zhang et al., 2021). The rate of delivery of functional CSCs into the PM of Arp2/3-inhibited cells (∼0.09 particles/µm^2^/min) was significantly reduced compared to mock-treated wild-type cells (0.12 particles/µm^2^/min; Fig. 1F). These results suggest that the Arp2/3 complex contributes to the secretion of CSCs to the PM, but does not affect the intracellular trafficking that brings SmaCCs from the subcortical region to the cortical region.

### The Arp2/3 complex participates in the late stages of CSC exocytosis, especially vesicle tethering

To evaluate whether the Arp2/3 complex is involved in the later stages of CSC exocytosis, we measured the tethering, docking, and fusion of CSC exocytic vesicles at the PM by tracking individual CSCs based on the previously established single-particle insertion assay (Fig. 2, A and B; Gutierrez et al., 2009; Zhang et al., 2019; Zhang et al., 2021; Zhang and Staiger, 2021; Zhu and McFarlane, 2022). The average particle pause time was 90 ± 34 s in the mock-treated wild type, and a shorter or longer pause time was defined as the mean value minus (< 56 s) or plus (> 124 s) 1 SD, respectively (Gutierrez et al., 2009; Zhang et al., 2019; Zhang et al., 2021; Zhang and Staiger, 2021). We categorized insertion events based on their pause time (representing tethering and docking) and whether they were successfully inserted as evidenced by motility in the plane of the PM (Fig. 2B). The *arp2-2*, *arpc2*, and CK-666-treated cells all had significantly lower successful insertion rates (76, 77, and 78%, respectively) compared to the mock-treated wild-type cells (86%; Fig. 2C). Most CSC particles in mock-treated wild-type cells paused for 60–100 s, whereas Arp2/3-inhibited cells had more CSCs with a pause time greater than 120 s (Fig. 2D). The combination of decreased successful insertions at the PM as well as the prolonged CSC pause time in genetically or chemically inhibited cells indicate that the Arp2/3 complex is involved in the late stages of CSC exocytosis.

**Figure 2.**
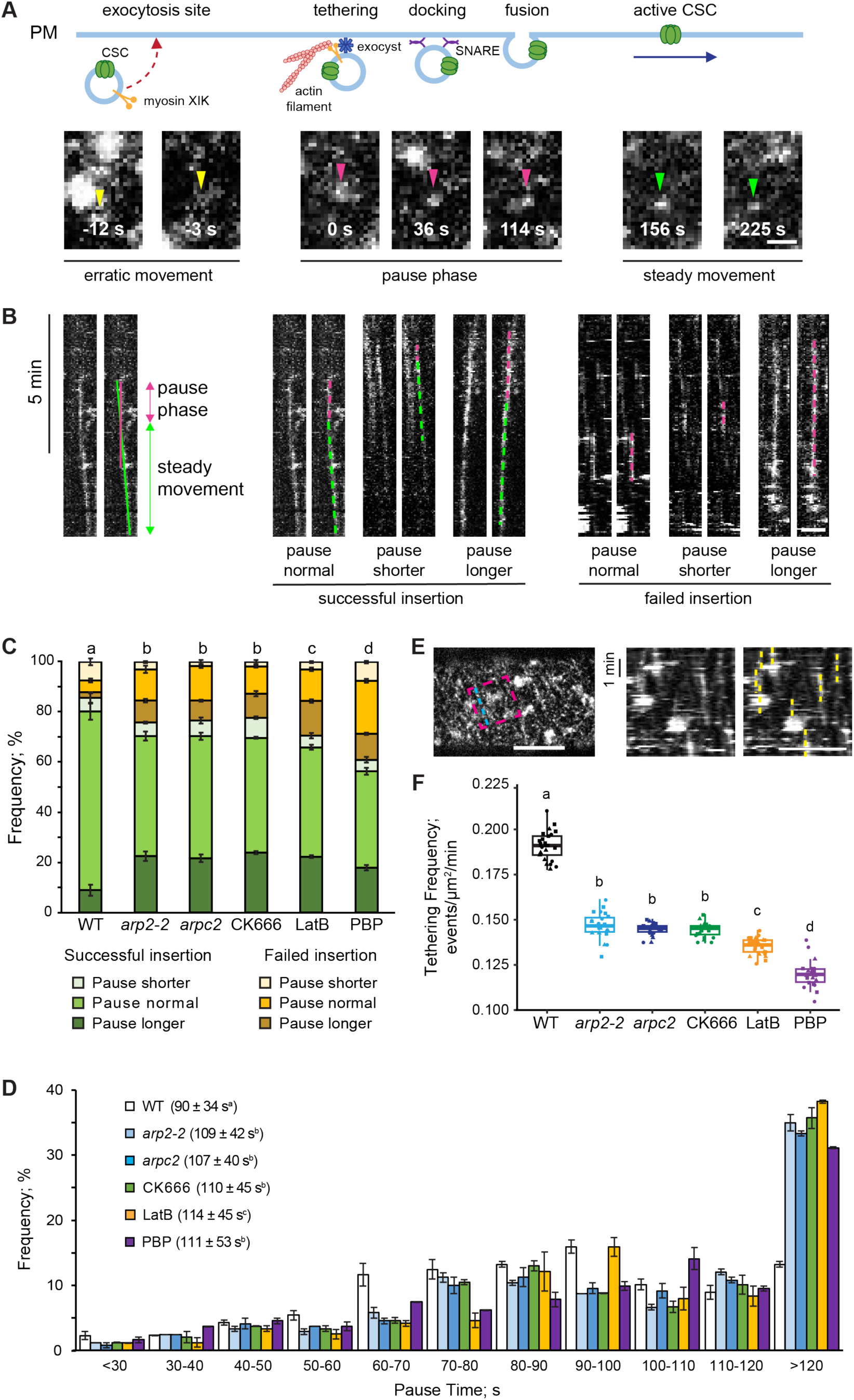
The Arp2/3 complex contributes to the late stages of CSC exocytosis. A, Representative time-lapse images of a typical CSC insertion event at the PM. A CSC particle (yellow arrowhead) arrived at the cortex and displayed an initial phase of erratic movement, which likely represents a vesicle delivered to the exocytosis site. The particle (magenta arrowhead) then remained stationary or paused for > 80 s, which likely represents tethering, docking, and fusion of the vesicle to the PM. After the CSC particle (green arrowhead) was inserted into the PM and potentially activated, it began a steady linear movement phase. Bar = 1 μm. B, Representative kymographs of the standard (left) and six subclasses (right) of CSC insertion event illustrating both the pause and steady movement phases, as well as successful and failed insertion events. The vertical line of best fit (magenta) represents the pause phase, whereas the sloped line of best fit along the moving trajectory (green) indicates the particle’s steady movement. The intersection of these two lines was determined as the end of the pause phase. Bar = 2 μm. C, The proportion of six types of insertion events described in (B) for mock-treated wild-type (WT), *arp2-2*, *arpc2*, CK-666-, LatB-, and PBP-treated cells, respectively. A shorter or longer pause time was defined as the mean value (90 ± 34 s) of particle pause time in mock-treated wild-type cells minus (< 56 s) or plus (> 124 s) 1 SD, respectively. D, Distribution of CSC pause times for insertion events measured in (C). For C and D, values shown are mean ± SE from three independent experiments (for each experiment, at least 80 CSC insertion events were measured from at least 7 seedlings per genotype/treatment). Letters [a–d] denote genotypes or treatments that show statistically significant differences with other groups by Chi-square test (P < 0.05). E, Representative single-frame image of a hypocotyl epidermal cell shows CSCs at the PM focal plane. An ROI (magenta dashed box) parallel to the CSC tracks (blue dashed line) was selected to extract all kymographs. A representative kymograph generated from the marked tracks shows the CSC trajectories at the PM and vertical yellow dashed lines indicate the pause phase and were counted as tethering events if they lasted longer than 30 s. Bars = 5 μm. F, Quantitative analysis of the frequency of CSC vesicle tethering. The frequency of vesicle tethering was significantly reduced in *arp2-2*, *arpc2*, and CK-666-treated cells when compared to that in mock-treated wild-type cells, but was not as severely decreased when compared to that in PBP- or LatB-treated cells. In the box-and-whisker plot, boxes show the interquartile range (IQR) and the median, and whiskers show the maximum-minimum interval of three biological repeats with independent populations of plants. The three independent experiments are represented with different symbols. n ≥ 21 seedlings (at least 7 seedlings were measured per independent experiment). Letters [a–d] denote genotypes or treatments that show statistically significant differences with other groups by one-way ANOVA with Tukey’s post-hoc test (P < 0.05).

To further investigate what specific step(s) the Arp2/3 complex participates in during exocytosis, we measured the frequency of CSC vesicle tethering events, as described previously (Fig. 2E; Zhang and Staiger, 2021). Here, we found that inhibition of the Arp2/3 complex led to a significantly lower tethering frequency compared to that in mock-treated wild-type cells (Fig. 2F). These results suggest that, during the late stages of CSC exocytosis, the Arp2/3 complex participates in the delivery of functional CSCs to the PM by facilitating the tethering and/or fusion of exocytic vesicles to the PM. This is similar to the previously described role for actin, but less pronounced than the role of myosin XI, during the late stages of exocytosis (Sampathkumar et al., 2013; Zhang et al., 2019; Zhang et al., 2021; Zhang and Staiger, 2021).

### CSCs pause transiently before internalization from the PM

When actin or the Arp2/3 complex were genetically or chemically inhibited, the density of PM-localized CSCs remained unchanged, even though secretion of CSCs to the PM was significantly reduced (Fig. 1; Fig. 2; Sampathkumar et al., 2013; Zhang et al., 2019). One explanation for these contradictory results is that the internalization of CSCs from the PM might also be disrupted under these conditions. To date, no study directly visualizes and quantifies specific steps of CSC internalization events at single-particle resolution, as has been done extensively for CSC exocytosis (Zhu and McFarlane, 2022; McFarlane, 2023). This is perhaps due to the crowded nature of CSCs in the PM of growing epidermal cells as well as their lifetime of 12–13 min during cellulose production, making high spatiotemporal resolution imaging of individual CSC internalization events technically challenging.

To aid in the detection of single CSC internalization events and to investigate the molecular mechanisms, we used mEOS2, a photoconvertible fluorescent protein (McKinney et al., 2009), to construct a new fluorescent-tagged CESA6 reporter and selected a transgenic complementation T3 line that rescued the growth phenotype of the *cesa6/prc1-1* mutant (Supplemental Fig. S3). Similar to a conventional YFP-CESA6-marked line, the mEOS2-CESA6 expressing line in the *prc1-1* background showed similar hypocotyl growth (Supplemental Fig. S4, A and B), cellulose content (Supplemental Fig. S4C), and *CESA6* expression level (Supplemental Fig. S4D) compared to wild type. In addition, cells expressing mEOS2-CESA6 or YFP-CESA6 showed similar distribution pattern and abundance (Supplemental Fig. S5, B and D) as well as motility (Supplemental Fig. S5, A and C) in the PM. These results validate that mEOS2-CESA6 is a functional fusion protein that can be used to visualize the dynamic activities of CESAs or CSCs in living cells. For mEOS2-CESA6 photoconversion, a region of interest (ROI) with a size of 80 x 80 pixels (113.2 µm^2^) was selected and irradiated with 405-nm laser light, which efficiently switched the emission of PM-localized mEOS2-CESA6 from green to red fluorescence (Supplemental Fig. S6). This photoconversion allowed the separation of cytoplasmic CESAs and PM-localized CSCs, facilitating the tracking of CSC dynamics at the PM, especially during CSC internalization. It is important to note that irradiation with the 405-nm laser for photoconversion did not alter the motility of mEOS2-CESA6 in the PM (Supplemental Fig. S5, A and C).

For single-particle resolution of CSCs and establishment of an internalization assay, seedlings were directly mounted in either mock or inhibitor solutions, photoconversion was excuted, and time-lapse movies were collected at 2-s intervals for at least 8 min with SDCM. Similar to the CSC insertion assay (Fig. 2; Gutierrez et al., 2009; Zhang et al., 2019), any CSC that exhibited steady movement after photoconversion was considered an active CSC, and an active CSC that completely disappeared from the PM focal plane was counted as a CSC internalization event (Fig. 3A). CSCs have been shown to be internalized by clathrin-mediated endocytosis (CME), and live-cell tracking of CME machinery proteins shows that these particles often display a pause phase lasting tens of seconds prior to their disappearance from the PM, likely representing coat initiation and assembly as well as the membrane invagination process during endocytosis (Bashline et al., 2013; Bashline et al., 2015; Sánchez-Rodríguez et al., 2018). Indeed, we observed a total of 2244 CSC internalization events and 74% of the events exhibited a pause phase, whereas 26% of the events did not show a visible pause phase (Fig. 3, B and F). To further quantify CSC internalization, we found that the events could be categorized into four subclasses based on the duration of the pause phase: (1) particles pausing for 40–70 s; (2) particles with a significantly shorter pause phase; (3) particles with a significantly longer pause; and (4) particles that disappeared without any pause phase (Fig. 3B). In mock-treated wild-type cells, the average pause time for CSC internalization events was 54 ± 21 s. Any pause time < 33 s (mean minus 1 SD) or > 75 s (mean plus 1 SD) was considered shorter (16% of total events) or longer (13%) than normal (71%), respectively (Fig. 3, D and E).

**Figure 3.**
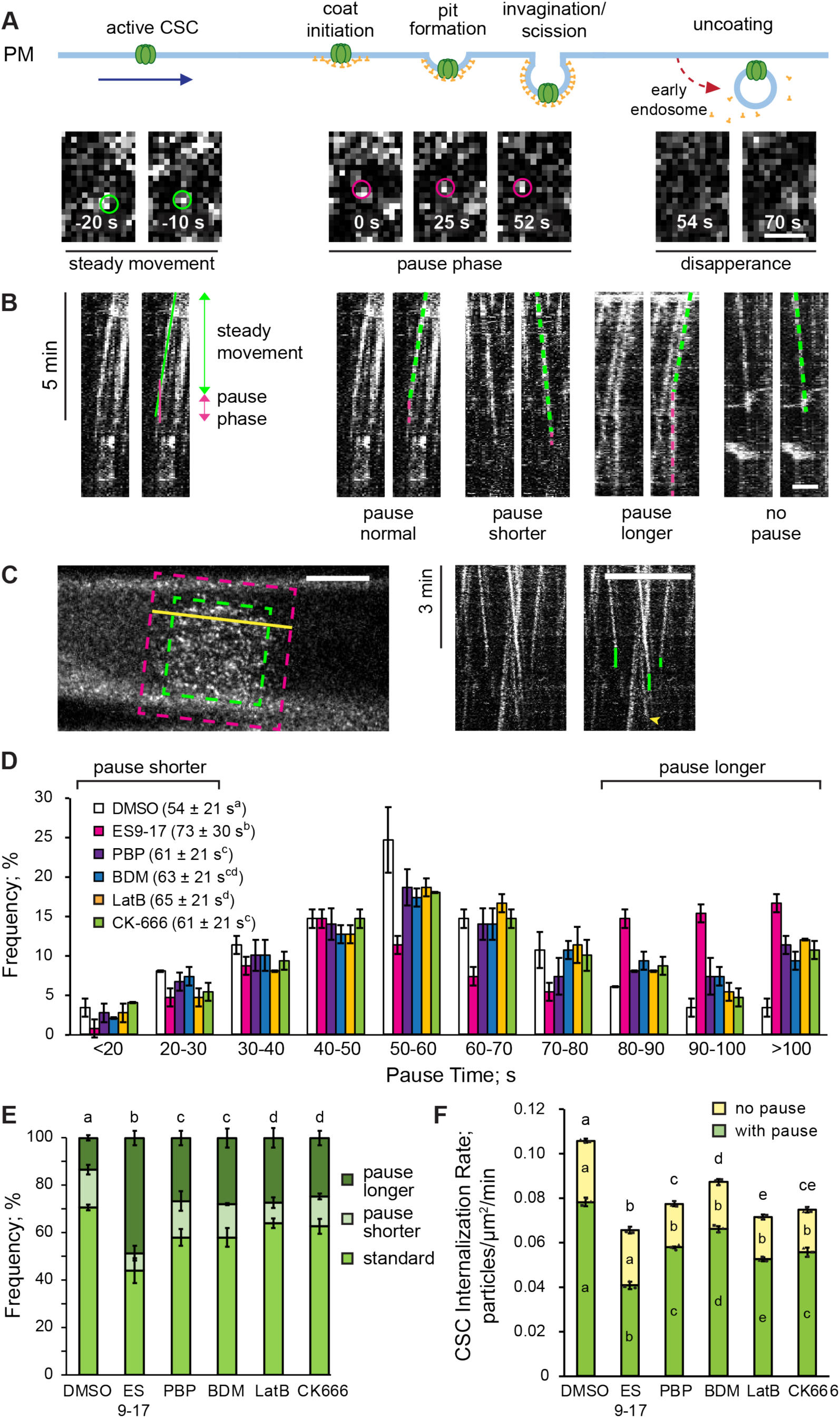
The actin cytoskeleton and myosin are involved in CSC internalization. A, Representative images of a CSC internalization event at the PM in a hypocotyl epidermal cell expressing mEOS2-CESA6 and following photoconversion from green to red (images shown are from the red channel). An active CSC (green circles) moved with a steady linear motion in the PM. Subsequently, the CSC (magenta circles) stopped moving and remained stationary for 50–60 s, which presumably represents the recruitment of coat components, coat assembly, membrane invagination, and the scission of an endocytic vesicle. After the CSC endocytic vesicle was successfully separated from the PM, it disappeared from the cortical region. Bar = 1 μm. B, Representative kymographs of the standard (left) and four subclasses (right) of CSC internalization event show both the steady movement and the pause phase. The sloped line of best fit (green) aligns with the moving trajectory representing the particle’s steady movement in the PM, whereas the vertical line of best fit (magenta) indicates the pause phase during the internalization of a CSC. The intersection of these two lines represents the beginning of the pause phase. Bar = 2 μm. C, Representative single-frame image of a hypocotyl epidermal cell expressing mEOS2-CESA6 shows CSCs at the PM focal plane. An ROI (green dashed box) on the PM was treated with laser photo-conversion, and a larger ROI (magenta dashed box) parallel to the CSC movement tracks (yellow solid line) was selected to extract all kymographs in the red channel. A representative kymograph generated from the marked CSC movement track (yellow solid line) shows the CSC trajectories at the PM. Vertical lines (green) indicate the pause phase and were counted as CSC internalization events with a pause phase. CSC trajectories that disappeared without a vertical line (yellow arrowhead) were counted as a CSC internalization event without a pause phase. Bar = 5 μm. D, Distribution of particle pause times for CSC internalization events measured from cells treated with mock (0.5% DMSO) solution, 70 μM ES9-17 (a clathrin inhibitor), 10 μM PBP, 30 mM BDM, 10 μM LatB, or 10 μM CK-666 for 30 min prior to imaging. The average pause times for internalization events in inhibitor-treated cells were significantly increased compared to mock-treated cells, especially with ES9-17 treatment. E, The proportion of three subclasses of internalization event described in (B) and measured in (D) were plotted. A shorter or longer pause time was defined as the mean value (54 ± 21 s) of particle pause time in mock-treated wild-type cells minus (< 33 s) or plus (> 75 s) 1 SD, respectively. The proportion of longer pause events was significantly increased in inhibitor- treated cells compared to mock-treated ones. For D and E, values are means ± SD from three independent experiments (for each experiment, at least 50 events were measured from at least 7 seedlings per treatment). For average pause time, letters [a–c] denote treatments that show statistically significant differences with other treatments by one-way ANOVA with Tukey’s post-hoc test (P < 0.05). For the distribution of the CSC particle pause time and the proportion of three subclasses of CSC internalization events, letters [a–d] denote treatments that show statistically significant differences with other treatments by Chi-square test (P < 0.05). F, Quantitative analysis of the CSC internalization rate as determined by photoconversion experiments as depicted in (C). The overall rate of CSC internalization was significantly reduced in inhibitor-treated cells compared to mock-treated cells, especially following ES9-17 treatment. The proportion of CSC internalization events with a pause phase (green bars) was significantly reduced in all inhibitor-treated cells compared to mock-treated cells. The proportion of CSC internalization events without a pause phase (yellow bars) was significantly reduced in PBP-, BDM-, LatB-, and CK-666-treated cells compared to mock- or ES9-17-treated cells. Values are means ± SE from three independent experiments (for each experiment, at least 7 seedlings per treatment were measured). Letters [a–e] denote treatments that show statistically significant differences with other treatments by one-way ANOVA with Tukey’s post-hoc test (P < 0.05).

If the pause phase is associated with clathrin pit formation, membrane invagination and scission, we expect that disruption of CME will alter the duration of the pause phase and subsequent internalization of CSCs. To test this, we treated hypocotyl epidermal cells with ES9- 17, a small molecule inhibitor that targets clathrin heavy chain (Dejonghe et al., 2019), and found that the average particle pause time (73 ± 30 s) as well as the proportion of long pause events (48.7%) in ES9-17-treated cells were significantly increased compared to mock-treated cells (Fig. 3, D and E). These results suggest that the CSC pause phase prior to internalization correlates with CME.

To further confirm the involvement of CME in CSC internalization, we revised the CSC tethering assay to quantitatively assess CSC internalization rate. The same time-lapse movies from the single-particle assay discussed above were used for analysis of internalization rate by kymographs. To calculate the number of internalization events within a unit area over a set duration of time, a larger ROI with a size of 100 x 100 pixels (176.9 µm^2^) was selected after photoconversion to encompass a majority of CSCs that migrated outside of the original 80 x 80 pixels (113.2 µm^2^) ROI over a 6-min time span (Fig. 3C). All internalization events within the larger ROI during the first 6 minutes were tracked by analysis of kymographs. In this assay, any moving trajectory (a line with a slope on the kymograph) that reached an endpoint or disappeared was counted as a CSC internalization event. For these events, moving trajectories ending with a vertical line were considered internalization events with a pause phase, regardless of the duration, whereas trajectories ending without a vertical line were counted as internalization events without a pause phase (Fig. 3C). In mock-treated wild-type cells, the average internalization rate was 0.106 ± 0.006 events/µm^2^/min (Fig. 3F), which is slightly lower than the frequency of CSC secretion (0.12 particles/µm^2^/min; Fig. 1F). This is probably because we did not include CSC particles that moved faster than average and migrated out of the 100 x 100 pixels ROI and were internalized outside of the ROI.

Acute treatment with 70 µM ES9-17 caused a 38% reduction in the total internalization rate (0.066 ± 0.005 events/µm^2^/min); however, the frequency of internalization events without a pause phase was only decreased by ∼10% (Fig. 3F). Collectively, these results suggest that CME is one of the major CSC internalization pathways, and the duration of the pause phase of a CSC particle prior to internalization is a key feature of this process.

### Actin and myosin are involved in CSC internalization

The involvement of actin and myosin in the internalization of plant plasma membrane proteins, especially CME, remains controversial (Zhang et al., 2019; Narasimhan et al., 2020; Diao and Huang, 2021; Aniento et al., 2022). To elucidate whether actin and myosin participate in CSC internalization, we disrupted actin assembly with LatB or CK666 and inhibited myosin activity with pentabromopseudilin (PBP; Zhang et al., 2019) or 2,3-butanedione monoxime (BDM; Zhang et al., 2019). Both actin (61–65 s) and myosin inhibition (61–63 s) caused significantly prolonged pause times compared to mock treatment (Fig. 3, D and E). Additionally, for the CSC internalization rate (Fig. 3F), there was a 27% reduction after PBP treatment (0.077 ± 0.006 particles/µm^2^/min), a 17% reduction after BDM treatment (0.087 particles/µm^2^/min), a 32% reduction after LatB treatment (0.072 ± 0.006 particles/µm^2^/min), and a 29% reduction with CK- 666 treatment (0.075 ± 0.005 particles/µm^2^/min). Notably, compared to ES9-17 treatment, both actin and myosin inhibition led to a 23–31% reduction in internalization rate without a pause phase (Fig. 3F). These results suggest that both actin and myosin are involved in CSC internalization.

### The Arp2/3 complex also contributes to CSC internalization

Since acute treatment with CK-666 implicates the Arp2/3 complex in CSC internalization (Fig. 3, D–F), we further tested this by analyzing single-particle internalization events for CSCs in *arp2/3* mutants. To do this, we crossed the *arp2-1* and *arpc2* mutants with a line expressing mEOS2- CESA6 in the *prc1-1* background and recovered both *arp2-1* and *arpc2* homozygous mutants, as well as wild-type sibling lines, expressing mEOS2-CESA6 in the *prc1-1* background. We measured CSC particle pause time as well as internalization rate in *arp2-1*, *arpc2*, and CK-666- treated cells. We found that both genetic and chemical inhibition of the Arp2/3 complex significantly prolonged the pause phase of CSC particles during internalization and elevated the proportion of long pause events (Fig. 4, A and B). Furthermore, the CSC internalization rate, with or without a pause phase, was significantly decreased in Arp2/3-inhibited cells (∼25–30% reduction; Fig. 4C). Additionally, there were no significant differences between CK-666-treated wild-type cells and mock-treated *arp2/3* mutants. Collectively, these results indicate that the Arp2/3 complex is not only involved in the late stages of CSC exocytosis but also plays an important role in CSC internalization, which is critical for cells to keep a homeostatic level of active CSCs in the PM.

**Figure 4.**
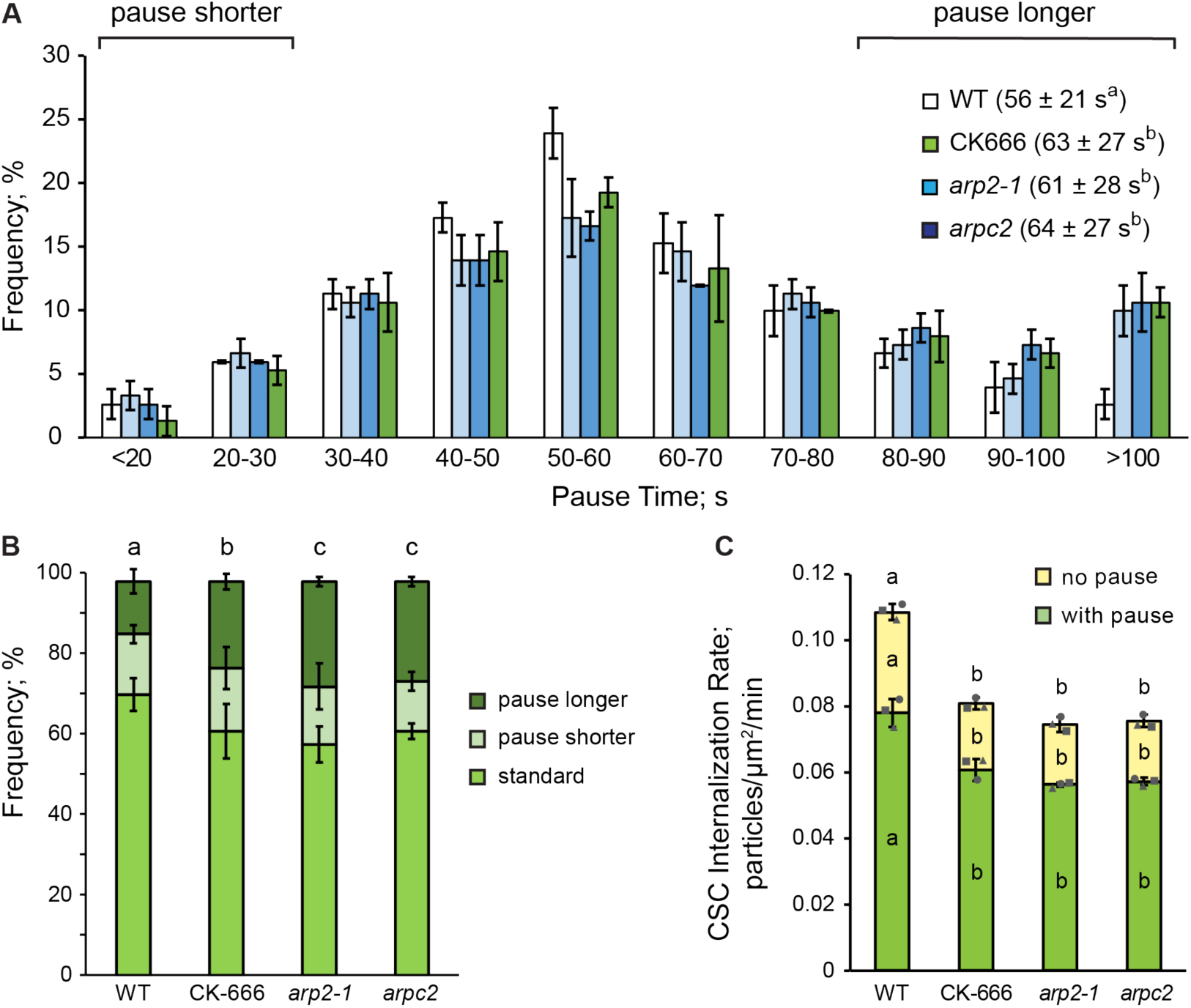
The Arp2/3 complex contributes to the rate of CSC internalization. A, Distribution of particle pause times for CSC insertion events measured from wild-type (WT), *arp2-1*, or *arpc2* cells treated with mock (0.5% DMSO) solution or from WT treated with 10 μM CK-666 for 5 min prior to imaging. The average pause times for internalization events in *arp2-1*, *arpc2*, and CK-666-treated cells were all significantly increased compared to mock-treated wild-type cells. Values are means ± SD from three independent experiments (for each experiment, at least 50 events from at least 7 seedlings per treatment were measured). Letters [a–b] denote genotypes or treatments that show statistically significant differences with other groups by one-way ANOVA with Tukey’s post-hoc test (P < 0.05). B, The proportion of three subclasses of internalization events as described in Figure 3 and measured in (A). Shorter or longer pause times were defined as the mean value (56 ± 21 s) of particle pause time in mock-treated wild-type cells minus (< 35 s) or plus (> 78 s) 1 SD, respectively. The proportion of pause longer events was significantly increased in Arp2/3-inhibited compared to mock-treated cells. Values are means ± SD from three independent experiments (for each experiment, at least 50 events from at least 7 seedlings per treatment were measured). Letters [a–c] denote genotypes or treatments that show statistically significant differences with other groups by Chi-square test (P < 0.05). C, Quantitative analysis of CSC internalization rate. The rate of CSC internalization was significantly reduced in cells without functional Arp2/3 complex when compared to mock-treated cells. The proportions of CSC internalization with or without a pause phase were both significantly reduced in Arp2/3-inhibited cells when compared to mock-treated cells. Values are means ± SD from three independent experiments (for each experiment, at least 7 seedlings per treatment were measured). Letters [a–b] denote genotypes or treatments that show statistically significant differences with other groups by one-way ANOVA with Tukey’s post-hoc test (P < 0.05).

## Discussion

In this study, we focused on investigating the involvement of actomyosin and the Arp2/3 complex in plant exo- and endocytic trafficking. We constructed a functional photoconvertible mEOS2-CESA6 reporter and developed new quantitative live-cell imaging approaches to directly visualize and assess the CSC dynamics at single-particle resolution. This mEOS2- CESA6 reporter allowed us to observe and quantitatively measure individual CSC internalization events by isolating and tracking active PM-localized CSCs in the red channel after photoconversion. In wild-type Arabidopsis hypocotyl epidermal cells, we found that a majority of CSCs exhibited a pause or static phase prior to internalization and that the rest of CSCs disappeared from the PM directly without evidence of pausing. These two subpopulations of CSC internalization events suggest the existence of multiple CSC internalization pathways.

Using acute treatment with the clathrin heavy chain inhibitor, ES9-17 (Dejonghe et al., 2019), we confirmed that CME is a key pathway for CSC internalization. Inhibition of CME resulted in a significantly prolonged pause time as well as a dramatic drop in the internalization rate. These results are consistent with previous findings that demonstrate several CME participants, including μ2 subunit of AP2 complex, TWD40-2, TPLATE and TML subunits of TPLATE COMPLEX, as well as dynamin-related proteins, show a pause phase on the PM, and their mutants lead to abnormal CSC distribution and slower CSC motility on the PM (Bashline et al., 2013; Bashline et al., 2015; Sánchez-Rodríguez et al., 2018). However, CME is a multi-step process that requires the cooperation of multiple different players during specific stages (Gnyliukh et al., 2024). Further co-localization analysis of individual CME participants and CSC particles during internalization, especially the pause phase, as well as quantitatively describing CSC internalization events in these CME mutants will be necessary to establish a correlation between CME and CSC pause phase during internalization and to better understand the molecular mechanisms of CSC internalization via CME.

Different studies make conflicting conclusions about the involvement of actin or myosin in FM4-64 or FLS2 endocytosis (Robatzek et al., 2006; Beck et al., 2012; Moscatelli et al., 2012; Sampathkumar et al., 2013; Kang et al., 2014; Mao et al., 2016; Zhang et al., 2019; Zhang et al., 2019; Narasimhan et al., 2020; Diao and Huang, 2021; Aniento et al., 2022). The recent study visualizing the clathrin coat/basket in plant cells with scanning electron microscopy showed the absence of a dense actin network around CCPs, suggesting that actin does not participate in CCP formation or membrane invagination (Narasimhan et al., 2020); however, the same study also revealed that actin is required for translocating CCVs away from the PM and guiding EE/TGN movements, which indicates an important role of actin in directing and organizing early post-endocytic trafficking during internalization. Our results demonstrate directly that the actin-myosin network contributes to CSC internalization. It is possible that actomyosin is involved in the internalization of large multiprotein complexes, such as CSCs.

Inhibition of actin led to a ∼32% reduction in the internalization rate of CSC particles, whereas the CSC delivery rate was reduced by ∼40%. This roughly equivalent reduction in both CSC secretion and internalization could explain the unaltered density of PM-localized CSCs following LatB treatment or in an *act2/7* double mutant (Sampathkumar et al., 2013; Zhang et al., 2019). On the other hand, myosin inhibition resulted in a 17–26% reduction in CSC internalization rate but a 57% decrease in CSC delivery rate, which coincides with the ∼20% reduction in the density of CSCs at the PM (Fig. 1; Fig. 3) (Zhang et al., 2019; Zhang and Staiger, 2021). The additional function of myosin XIK interacting with the exocyst complex to facilitate vesicle tethering during exocytosis could explain the difference between CSC exo- and endocytosis with myosin inhibition. In addition, inhibition of actin or myosin led to a significant reduction in the frequency of CSC internalization events, with and without a pause phase, suggesting that actomyosin participates in multiple CSC internalization pathways regardless of the requirement of a pause phase prior to internalization.

Our results also reveal a new function for the Arp2/3 complex in plant cells, which participates in both the secretion and internalization of CSCs. Genetic or chemical inhibition of the Arp2/3 complex caused a similar phenotype with actin disruption, including unchanged CSC density and reduced CSC secretion and internalization. This similarity suggests that the Arp2/3 complex contributes to exo- and endocytic trafficking of CSCs, possibly through mediating the dynamic homeostasis of cortical actin (Xu et al., 2024), which is a unique function of the Arp2/3 complex that has never been described previously in plant cells.

## Materials and methods

### Plant materials and growth conditions

All Arabidopsis (*Arabidopsis thaliana*) plants used in this study were in Columbia-0 (*Col-0*) background. The *ARP2* T-DNA insertion mutant, *arp2-2* (SALK_077920), was obtained from the Arabidopsis Biological Resource Center (Ohio State University), the *ARPC2* point mutation line, *arpc2* (G1217A), was kindly provided by Dr. Daniel B. Szymanski (Purdue University), and the homozygous *prc1-1* mutant line expressing YFP-CESA6 was kindly provided by Dr. Ying Gu (Penn State University). Both *arp2-2* and *arpc2* mutant lines were crossed to the homozygous *prc1-1* mutant line expressing YFP-CESA6 (Paredez et al., 2006), and homozygous mutants as well as corresponding wild-type siblings in the presence of *prc1-1* were recovered from F2 populations.

Seeds were surface sterilized and stratified at 4°C for 3 days on half-strength Murashige and Skoog (MS) plates supplemented with 1% (w/v) sucrose and 1 % (w/v) agar. For dark-growth, seedlings were grown in continuous darkness after exposing the plates to light for 4 h. For light-growth, seeds were under long day conditions (16 h light/8 h dark) at 21°C provided by Philips F32T8/L941 Deluxe Cool White bulbs with a light intensity of 120–140 mmol m^-2^s^-1^.

### Drug treatments

For short-term live cell treatments, individual seedlings were mounted in the inhibitor solution for 5 min in the dark prior to each imaging session. CK-666 (Sigma-Aldrich; St. Louis, MO, USA), LatB (Calbiochem; San Diego, CA, USA), and PBP (Adipogen; San Diego, CA, USA) were dissolved in DMSO to prepare a 5 mM stock solution. ES9-17 (Biosynth; Louisville, KY, USA) was dissolved in DMSO to prepare a 50 mM stock solution. BDM (Sigma-Aldrich) was dissolved in water immediately before use. FM4-64 (Invitrogen; Eugene, OR, USA) was dissolved in DMSO and used as a 20 μM solution.

### Cloning and transformation of the *mEOS2-CESA6* construct

The pUBN-CFP-DEST binary vector was used as a backbone to prepare the *mEOS2-CESA6* construct (Supplemental Fig. S3). Briefly, pUBN vector was digested with *Sac*I and *Sma*I to remove the ubiquitin promoter, CFP tag and Gateway related elements. The *CESA6* genomic sequence with terminator was amplified with primers F: AAAGAGCTCAAATCTAGAAAAGATATCATGAACACCGGTGGTCGGTT and R: AAACCCGGGGTGATCCACATCTTAAATATAT from wild type *Col-0* genomic DNA. The purified *CESA6* genomic sequence was digested with *Sac*I and *Sma*I, and ligated with same enzymes digested pUBN vector. The forward primer introduced *Xba*I and *Eco*RV restriction enzyme digestion sites for introducing CESA6 native promoter and mEOS2 sequence. The *CESA6* promoter (2.2 kb) was amplified from wild type *Col-0* genomic DNA with primers F: AAAGAGCTCTTTCTATTCTATAGTCTTGA and R: AAATCTAGAATTTGTCTGAAAACAGACACAG, and ligated with pUBN-CESA6 through digestion by *Sac*I and *Xba*I enzymes and ligation. mEOS2 was amplified from pRSETa-mEOS2 vector (Addgene; Watertown, MA, USA) with primers F: CTGTTTTCAGACAAATTCTAGAATGAGTGCGATTAAGCCAGA and R: CCACCGGTGTTCATGATATCTCGTCTGGCATTGTCAGGCA. pUBN-CESA6 with promoter was digested with *Xba*I and *Eco*RV enzymes and ligated with mEOS2 through Gibson assembly method. The *mEOS2-CESA6* construct was verified by wide-sequencing.

### RNA extraction and RT-qPCR

Total RNA was extracted from 6-day-old light-grown seedlings using TRIzol reagent (Invitrogen) according to the manufacturer’s protocol. RNA (0.5 µg) was reverse transcribed by the PrimeScript RT reagent kit with genomic DNA Eraser (Takara; Madison, WI, USA) using random and oligo-dT primer mix in a 10 µL reaction. Bio-Rad SsoAdvanced SYBR Green Supermix (Bio-Rad; Hercules, CA, USA) was used to perform the RT-qPCR in a LightCycler 96 thermocycler (Roche; Indianapolis, IN, USA). The relative gene expression levels were calculated using the 2−ΔΔCT method as described (Livak and Schmittgen, 2001) *AtCESA6* was amplified with primers (CESA6-F: ATGAACACCGGTGGTCGGTT; CESA6-R: CCATCAACAGTCAATTCGATCTC), *AtActin1* (ACTIN-F: GGTCACGACCAGCAAGATCAA, ACTIN-R: GGTCACGACCAGCAAGATCAA) was used as an internal control for gene expression normalization.

### Live-cell imaging

For all imaging acquisition, epidermal cells from the apical region of 3-d-old dark-grown hypocotyls were used unless otherwise stated. An Olympus IX-83 microscope mounted with a Yokogawa CSUX1-A1 spinning-disc confocal unit and an Andor iXon Ultra 897BV EMCCD camera were used to acquire snapshot and time-lapse images with an Olympus UPlanSApo oil objective (100X, 1.45 numerical aperture). YFP fluorescence was excited with a 514-nm laser line and emission collected through the 527–542-nm filter. CSC density, CSC motility and CSC insertion imaging as well as the cortical and subcortical YFP-CESA6 imaging were collected with MetaMorph software (version 7.8.8.0) as described previously (Zhang et al., 2019; Zhang et al., 2021). For FRAP and photoconversion experiments, images were collected with the same equipment and software mentioned above. Photobleaching of YFP-CESA6 was performed in a small, selected region (113.2 μm^2^) with an Andor Mosaic3 photomanipulation module and Andor iQ3 software (Andor Technology, Belfast, UK) with a 445-nm 1300 mW laser line at 35% power for 3 s, and time-lapse images were collected at the PM with a 5-s interval for 97 frames after photobleaching at the fourth/fifth frame. Photoconversion of mEOS2-CESA6 was performed in a selected region (113.2 μm^2^) using the same equipment as the photobleaching treatment with a 405-nm 1100 mW laser line at 25% power for 1 s, and time-lapse images were collected with a 2-s interval for 241 frames after photoconversion treatment.

### Image processing and quantitative analysis

All image processing and analyses were performed with Fiji Is Just Image J (Schindelin et al., 2012). Any image drift was corrected with the StackReg plug-in. All YFP-CESA6-related assays, including CSC density, motility, cortical and subcortical CESA compartment density, CSC delivery rate, single CSC insertion and CSC particle tethering rate were performed as described previously (Zhang et al., 2019; Zhang et al., 2021; Zhang and Staiger, 2021).

For CSC delivery rate analysis, newly-delivered CSCs from a smaller subregion (63.7 μm^2^) within the photobleached region during the first 5 min of recovery were manually counted, and only particles that exhibited a linear steady movement at the PM were counted as new delivery events (Gutierrez et al., 2009). For cortical and subcortical CESA compartment density analysis, only particles found in the first frame that showed a continuous mobility in the following frames and also had a size that was smaller than a TGN or Golgi compartment were counted as small CESA-containing compartments (SmaCCs). Z-series with focal planes at 0.2–0.4 μm below the PM were used as the cortical cytoplasm region and ones at 0.6–1.0 μm were used as the subcortical region.

For the CSC insertion analysis, time-lapse images were collected in the focal plane of the PM with 3-s intervals for 8 min. Only particles that showed *de novo* appearance at the PM followed by a pause phase of more than 5 frames (>15 s) were counted as new insertion events. The duration of the pause phase was determined by analysis of kymographs based on the trajectories of new insertion events. The straight vertical line at the beginning of an insertion event on the kymograph was considered to be the pause phase, and the straight line with a slope following the pause phase was considered as the steady movement of an active CSC particle. For quantitative analysis, at least 10 random insertion events were analyzed in each hypocotyl for at least 7 seedlings per genotype or treatment.

For the CSC tethering frequency analysis, the same time-lapse series used for the CSC insertion analysis were used. To quantify the frequency of tethering events, time-lapse series were rotated to align most CESA trajectories with the vertical line that was used to generate kymographs, and all tethering events that lasted for at least 30 s within a ROI of 60 x 60 pixels (63.7 μm^2^) at the PM were counted during the first 6 min.

The CSC internalization analysis using mEOS2-CESA6 was modified from the previously established CSC insertion assay (Gutierrez et al., 2009; Zhang et al., 2019). A small ROI of 80 x 80 pixels (113.2 µm^2^) was selected in the green (488 nm) channel and treated with 405-nm laser irradiation at 25% laser power for 1000 ms, and time-lapse images were collected at the PM with a 2-s interval for 8 min in the red/orange (561 nm) channel after the photoconversion. All particles that showed steady movement at the PM (or a sloped line in the kymograph) were considered active CSCs, and any active CSC that completely disappeared from the PM focal plane was counted as an internalization event. The appearance and duration of a pause phase were determined by analysis of kymographs. Only a vertical line in the kymograph was considered as a pause event. To determine the exact duration of the pause phase, a straight line was fitted along the moving trajectory and another line along the pause phase on the kymograph, and the intersection of these two lines was defined as the start of the pause phase. For quantification analysis, in each biological repeat, at least 7 random internalization events showing a pause phase were analyzed in each hypocotyl for at least 7 seedlings per genotype or treatment.

The CSC internalization rate analysis was modified from the previously established CSC tethering rate analysis (Zhang and Staiger, 2021). The same SDCM time-lapse images after photoconversion used for the single-particle internalization assay were used. To estimate an internalization rate (number of events per unit area and time), a larger ROI of 100 x 100 pixels (176.9 µm^2^) than the photoconversion area was chosen and all internalization events were measured within the larger ROI during the first 6 min by kymograph analysis. Individual time-lapse series were rotated to align most CESA trajectories with the vertical line that was used to generate kymographs. All moving trajectories (sloped lines on kymographs) that disappeared were counted as CSC internalization events. Any trajectory with a vertical line immediately before the disappearance on the kymograph was counted as a CSC internalization event with a pause phase. Any trajectory that disappeared directly without a vertical line on the kymograph was counted as a CSC internalization event without a pause phase. The internalization rate was calculated as the number of internalization events per unit area divided by elapsed time.

### FM4-64 internalization assay

7-day-old etiolated wild-type hypocotyls were pre-soaked in mock (0.5% DMSO) or inhibitor solution for designated times followed by incubation in 20 μM of FM4-64 with or without inhibitor for 6 min in the dark. Images of epidermal cells in the apical region of the hypocotyl were collected with SDCM as described above using a 60 x 1.42 NA UPlanSApo oil objective (Olympus). The number of internalized FM4-64 cytoplasmic particles in individual cells was counted, and the density was calculated as the number of particles per 100 μm^2^.

### Cell wall determination

7-day-old dark-grown hypocotyls without seed coat were used for cellulose measurement. Seedlings were ground into a fine powder in liquid nitrogen, and the homogenate was washed twice with 80% ethanol, once with 100% ethanol, once with 1:1 (v/v) methanol:CHCl3, and once with acetone. The alcohol-insoluble cell wall material (CWM) was then dried in the fume hood for 2 days. Cellulose content was measured as modified from the Updegraff method (Updegraff, 1969; Foster et al., 2010). Briefly, 2–3 mg of CWM was hydrolyzed with trifluoroacetic acid (TFA) and acetic-nitric reagent (AN, also called Updegraff reagent; acetic acid:nitric acid:water = 8:1:2 [v/v/v]) to yield crystalline cellulose. Alternatively, 2–3 mg of CWM was treated with only AN reagent to yield total cellulose. The amounts of glucose in the insoluble fractions from both the TFA and AN hydrolysis were prepared with Anthrone reagent and measured using the colorimetric assay compared to cellulose standards (DuBois et al., 1956).

### Statistical analyses

One-way ANOVA with Tukey’s post-hoc tests were performed in (GraphPad Prism 10) to determine significance among different genotypes or treatments. Chi-square tests were performed for statistically comparing parametric distributions and P values were calculated in Excel 15.32. Any difference with a P value less than 0.05 was considered significantly different. Detailed statistical analysis data are shown in Supplemental Data Set 1.

### Accession numbers

Sequence data from this article can be found in The Arabidopsis Information Resource (TAIR) under the following accession numbers: ARP2, AT3G2700; ARPC2, AT1G30825; CESA6, AT5G64740.

## Acknowledgements

This paper is dedicated to the memory of Dr. Chunhua Zhang, an excellent scientist, a great friend, and an invaluable colleague, who passed away on May 15, 2021. We thank Hongbing Luo (Purdue University) for excellent care and maintenance of plant materials, Dan Szymanski (Purdue University) for providing the *arpc2* line, and Ying Gu (Penn State University) for providing the homozygous *cesa6/prc1-1* line expressing YFP-CESA6. We are also grateful to the Arabidopsis Biological Resource Center (Ohio State University) for supplying the *arp2-1* and *arp2-2* T-DNA insertion lines.

## Author contributions

L.X. acquired and analyzed the data, prepared the figures, and wrote and revised the manuscript; L.H. acquired and analyzed the data; W.Z. revised the manuscript; C.Z. conceptualized experiments and acquired funding; and C.J.S. conceptualized the experiments, acquired the funding, and analyzed the data, and wrote and revised the manuscript.

## Supplemental data

The following materials are available in the online version of this article.

**Supplementary Figure S1.** Genetic disruption of the Arp2/3 complex reduces total and crystalline cellulose levels in the hypocotyl cell wall.

**Supplementary Figure S2.** Genetic or chemical inhibition of the Arp2/3 complex causes a modest decrease in CSC motility in the plasma membrane.

**Supplementary Figure S3.** Schematic diagram showing the mEOS2-CESA6 construct.

**Supplementary Figure S4.** The isolation and characterization of mEOS2-CESA6 transgenic lines.

**Supplementary Figure S5.** Two different CESA markers, mEOS2-CESA6 and YFP-CESA6, show similar dynamic properties on the PM.

**Supplementary Figure S6.** mEOS2-CESA6 is a functional, photoconvertible marker for tracking PM-localized CSCs.

**Supplementary Figure S7.** The internalization of FM4-64 is deficient in Arp2/3-inhibited cells.

**Supplementary Data Set 1.** Statistical analyses results and parameters.

