## Supplemental Figures for "Actomyosin and the Arp2/3 Complex Are Involved in the Internalization of Cellulose Synthase Complexes"

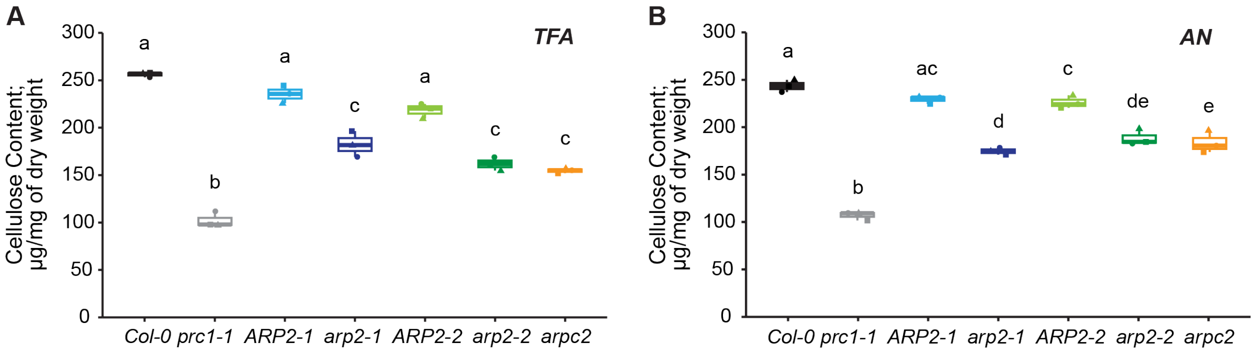


**Figure S1. Genetic disruption of the Arp2/3 complex reduces total and crystalline cellulose levels in the hypocotyl cell wall.**

The ethanol-insoluble portion of cell wall material (CWM) was extracted from 5-d-old etiolated hypocotyls of *cesa6/prc1-1*, *arp2-1*, *arp2-2*, and *arpc2* homozygous mutants as well as from the wild-type sibling lines *ARP2-1*, *ARP2-2*, and Col-0 respectively. A and B, The non-cellulosic component of CWM was hydrolyzed with 2 M trifluoroacetic acid (TFA; A) to determine the total cellulose content, or with acetic nitric reagent (AN; B) to determine the crystalline cellulose content. Total and crystalline cellulose content were significantly reduced in *prc1-1*, *arp2-1*, *arp2-2*, and *arpc2* mutants when compared to that in the wild-type lines. In box-and-whisker plots, boxes show the interquartile range (IQR) and the median, and whiskers show the maximum-minimum interval of three independent experiments. Letters [a–c] denote genotypes that show statistically significant differences with other genotypes by one-way ANOVA with Tukey's post-hoc test ($P < 0.05$).

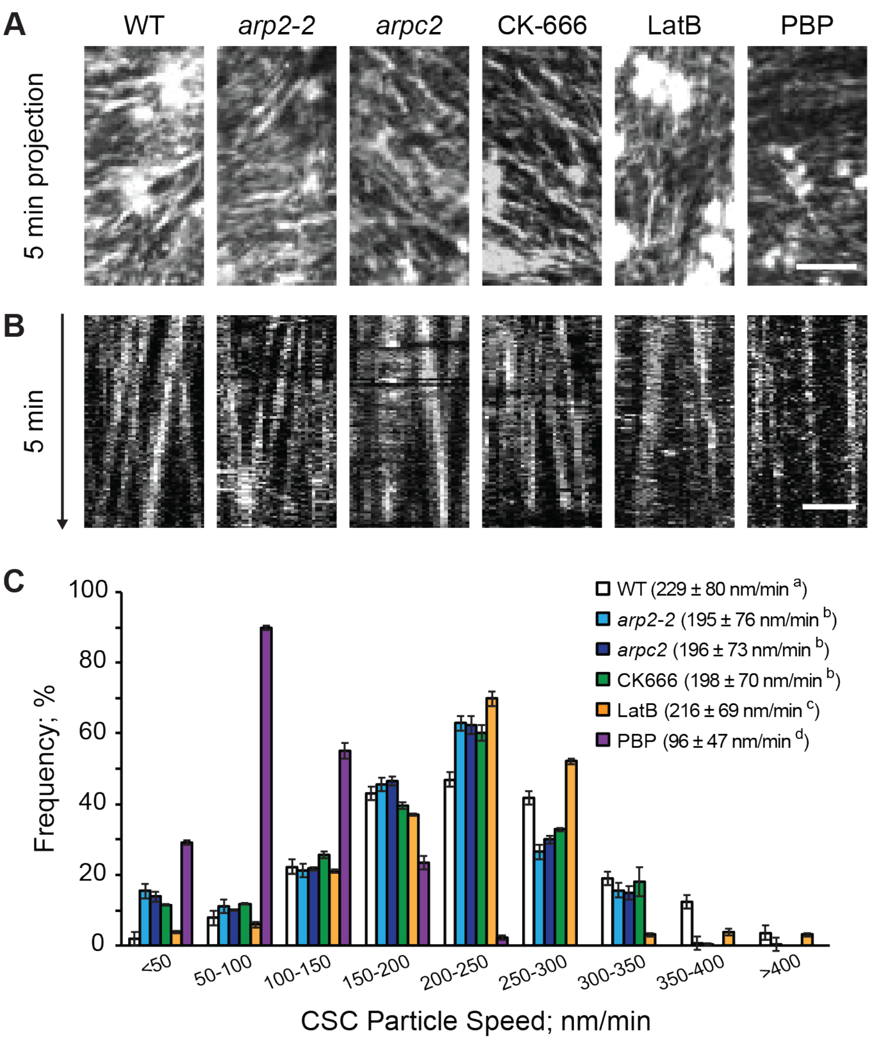


**Figure S2. Genetic or chemical inhibition of the Arp2/3 complex causes a modest decrease in CSC motility in the plasma membrane.**

A, Representative time projections of CSC trajectories in the PM. Time projections were generated from the average intensity of 161 frames at 3-s intervals collected from genotypes and chemical inhibitor treatments as described for Figure 1. Bar = 5 μm.
B, Representative kymographs of CSC particle movement at the PM during a 5-min interval. Bar = 2 μm.
C, Frequency distribution and mean value of CSC particle speed for wild-type, *arp2-2*, *arpc2* and chemical inhibitor treatments. Distributions for *arp2-2*, *arpc2*, or CK-666-treated wild-type cells were comparable to mock-treated wild-type cells, albeit the mean values were modestly, but significantly slower. In contrast, the reductions in mean speed and frequency distribution of CSCs in PBP-treated cells were markedly different when compared to mock-treated wild type. Values shown are mean ± SE from three independent experiments (for each experiment, at least 50 CSC trajectories were measured from at least 7 seedlings per genotype/treatment). Letters [a–d] denote genotypes/treatments that show statistically significant differences with other groups by one-way ANOVA with Tukey's post-hoc test ($P < 0.05$).

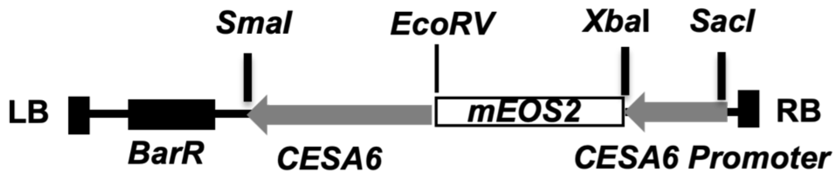


**Figure S3**. **Schematic diagram showing the mEOS2-CESA6 construct**.

LB, left border; RB, right border. BarR, Basta resistance gene.

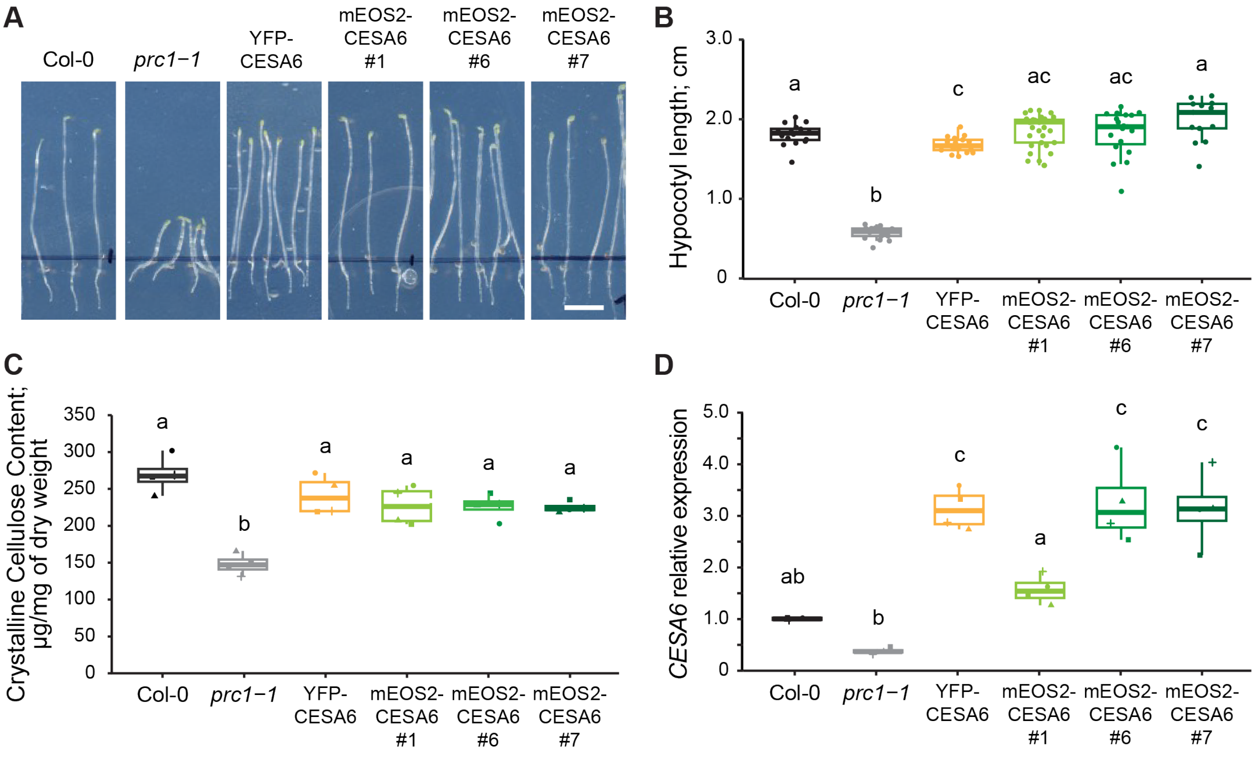


**Figure S4. The isolation and characterization of mEOS2-CESA6 transgenic lines**

A, Representative images of 5-d-old etiolated seedlings of wild type (Col-0), homozygous *prc1-1* (a null mutant for *cesa6*), and *prc1-1* complemented with *pCESA6::YFP-CESA6* or *pCESA6::mEOS2-CESA6*. Bar = 0.5 cm.

B, Seedlings expressing either YFP-CESA6 or three independent mEOS2-CESA6 complementation lines were able to fully rescue the reduced hypocotyl length of *prc1-1*. In box-and-whisker plots, boxes show the interquartile range (IQR) and the median, and whiskers show the maximum-minimum interval ($n \geq 10$ individual seedlings per genotype). Letters [a–b] denote genotypes that show statistically significant differences with other groups by one-way ANOVA with Tukey's post-hoc test ($P < 0.05$).

C, Both YFP-CESA6 and all three mEOS2-CESA6 complementation lines rescued the reduced crystalline cellulose content of *prc1-1*.

D, Transcript levels of *CESA6* in *prc1-1* seedlings expressing YFP-CESA6 or mEOS2-CESA6 (line #6 and #7) were comparable, albeit significantly higher than Col-0 wild type.

For box-and-whisker plots in in (C) and (D), boxes show the interquartile range (IQR) and the median, and whiskers show the maximum-minimum interval of four biological repeats with independent populations of plants ($n = 4$ independent experiments). Letters [a–c] denote groups that show statistically significant differences with other groups by one-way ANOVA with Tukey's post-hoc test ($P < 0.05$).

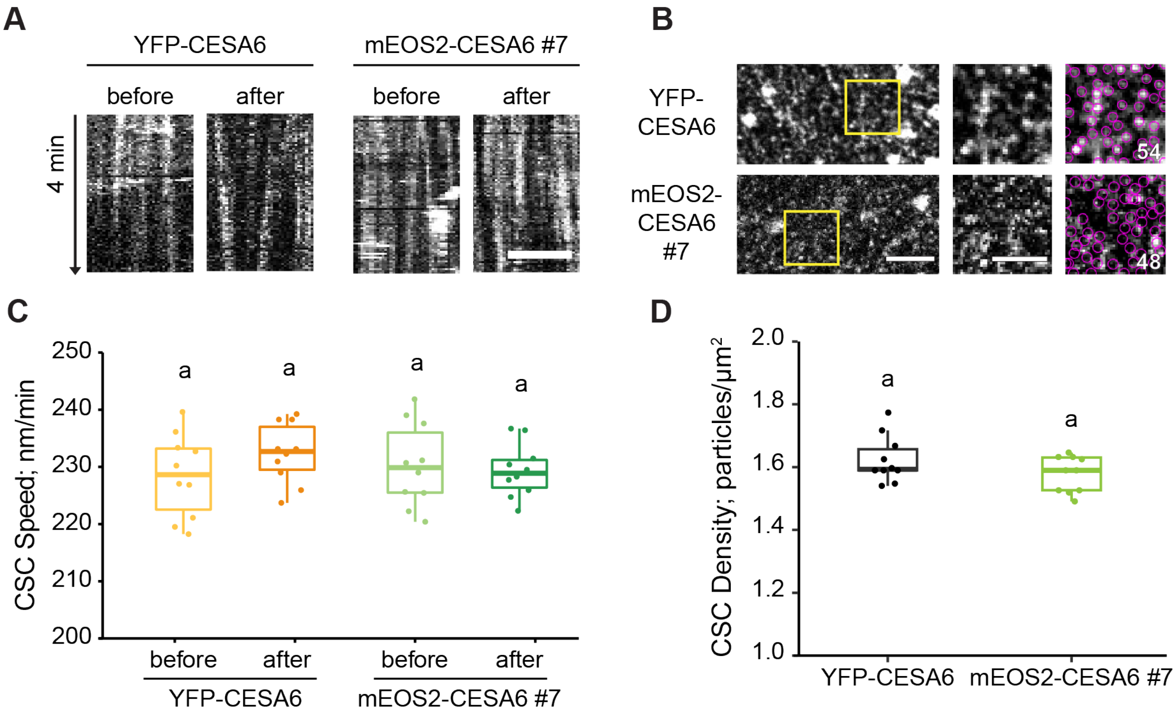


**Figure S5. Two different CESA markers, mEOS2-CESA6 and YFP-CESA6, show similar dynamic properties on the PM.**

A, Representative kymographs of CSC particle movement at the PM during a 8-min interval before or after photoconversion from green to red. Bar = 3 µm.
B, Representative single-frame images of epidermal cells from the apical region of 3-d-old etiolated hypocotyls expressing YFP-CESA6 or mEOS2-CESA6 collected by SDCM show the distribution of CSCs in the PM (left column). Bar = 5 µm. ROIs (boxes) from the left figures were magnified (middle column). Bar = 3 µm. The number of CSC particles detected by TrackMate in Image J from each ROI is shown in the right column (circles).
C and D, The average speed (C) or density (D) of CSCs at the PM did not show any significant difference between YFP-CESA6 and mEOS-CESA6. For box-and-whisker plots, boxes show the interquartile range (IQR) and the median, and whiskers show the maximum-minimum interval ($n \geq 10$ individual seedlings per genotype). Letters denote genotypes that show statistically significant differences with other groups by one-way ANOVA with Tukey's post-hoc test ($P < 0.05$).

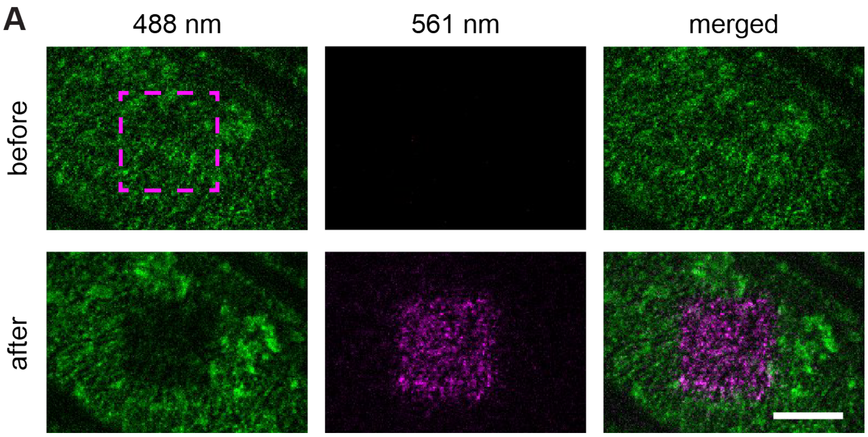


**Figure S6. mEOS2-CESA6 is a functional, photoconvertible marker for tracking PM-localized CSCs.**

A, Representative single-frame images of epidermal cells from the apical region of 3-d-old etiolated hypocotyls expressing mEOS2-CESA6 collected by SDCM show the distribution of CSCs on the PM. Bar = 5 µm. PM-localized mEOS2-CESA6 in an ROI of 80 X 80 pixel (113.2 $\mu m^2$; dashed box) was switched from green fluorescence (left column) to red (middle column) with 405-nm laser irradiation.

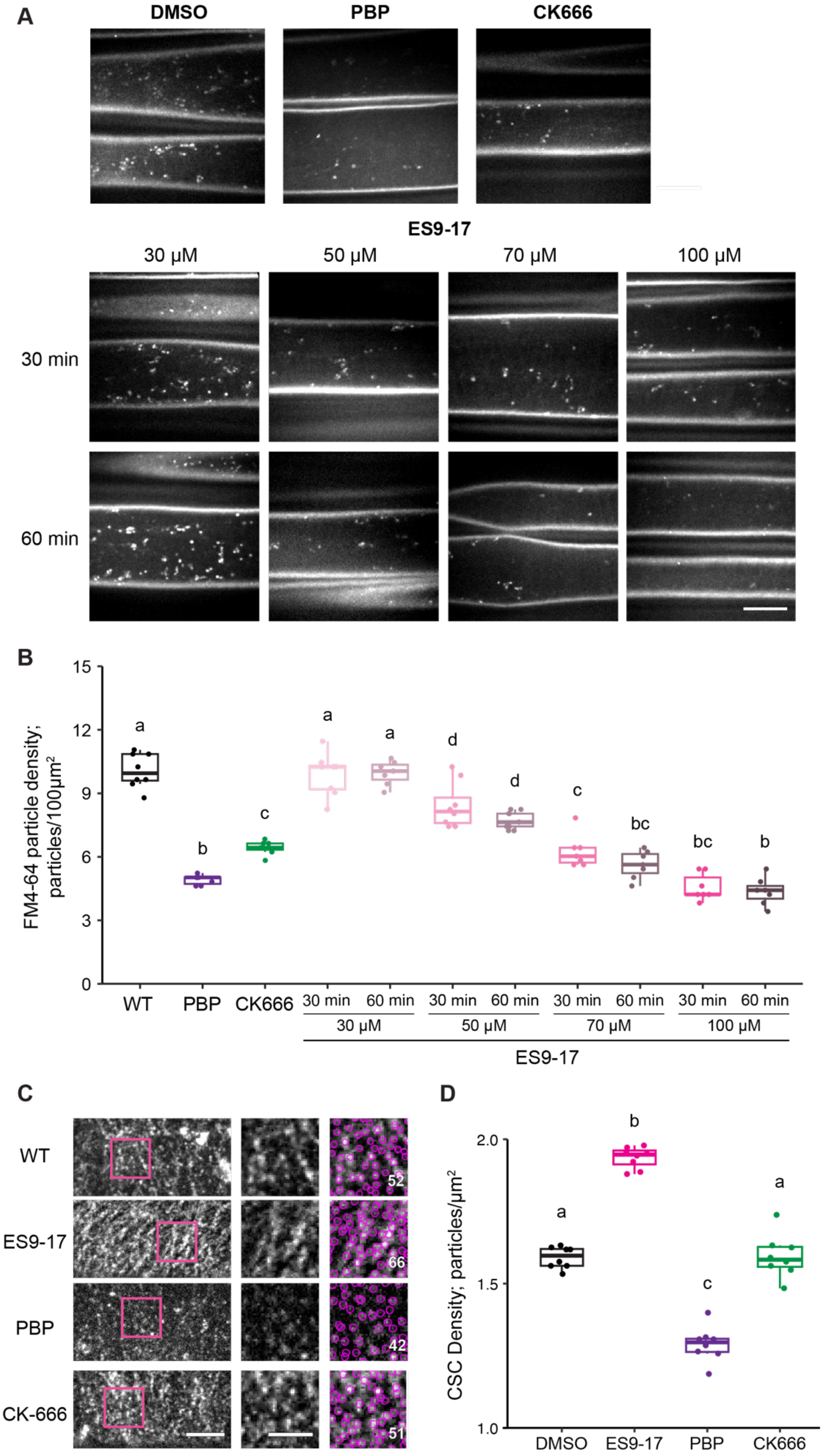
A
DMSO
PBP
CK666
ES9-17
30 µM
50 µM
70 µM
100 µM
30 min
60 min
B
FM4-64 particle density; particles/100µm²
15
12
9
6
3
0
a
b
c
a
a
d
d
c
bc
bc
b
WT
PBP
CK666
30 min
60 min
30 µM
50 µM
70 µM
100 µM
ES9-17
C
WT
ES9-17
PBP
CK-666
52
66
42
51
D
CSC Density; particles/µm²
2.0
1.5
1.0
a
b
c
a
DMSO
ES9-17
PBP
CK666

**Figure S7. The internalization of FM4-64 is deficient in Arp2/3-inhibited cells.**

A, Representative single-frame images show the internalization of FM4-64 in 3-d-old dark-grown hypocotyl epidermal cells. Seedings were pre-treated with mock (0.5% DMSO) solution for 60 min, 10 μM PBP for 15 min, or 10 μM CK666 for 15 min. Seedlings treated with ES9-17 were soaked in 30, 50, 70, or 100 μM ES9-17 solution for either 30 or 60 min. After inhibitor treatment, seedlings were treated with 20 μM FM4-64 for 5 min prior to imaging with SDCM. Bar = 5 μm.
B, Quantitative analysis of the density of FM4-64 marked cytoplasmic particles. The internalization of FM4-64 was significantly inhibited in PBP- or CK666-treated cells. Treatment with 70 μM ES9-17 for 30 min also reduced the FM4-64 internalization as significantly as PBP or CK666. In box-and-whisker plots, boxes show the interquartile range (IQR) and the median, and whiskers show the maximum-minimum interval ($n \geq 8$ individual seedlings per genotype/treatment). Letters [a–d] denote treatments that show statistically significant differences with other treatments by one-way ANOVA with Tukey's post-hoc test ($P < 0.05$).
C, Representative single-frame images of epidermal cells from the apical region of 3-d-old etiolated hypocotyls expressing YFP-CESA6 show the distribution of CSCs on the PM (left column). Bar = 5 μm. Magnified ROIs (magenta boxes) from figures are shown in the middle column. Bar = 3 μm.
D, Quantitative analysis of CSC density at the PM. The density of CSCs was significantly increased in ES9-17-treated cells and significantly reduced in PBP-treated cells compared to mock-treated cells, whereas CK-666-treated cells did not show any significant difference. In box-and-whisker plots, boxes show the interquartile range (IQR) and the median, and whiskers show the maximum-minimum interval of at least 7 seedlings per treatment. Letters [a–c] denote groups that show statistically significant differences with other treatments by one-way ANOVA with Tukey's post-hoc test ($P < 0.05$).
